# *Klebsiella pneumoniae* mutants resistant to ceftazidime/avibactam plus aztreonam, imipenem/relebactam, meropenem/vaborbactam and cefepime/taniborbactam

**DOI:** 10.1101/2021.11.21.469431

**Authors:** Naphat Satapoomin, Punyawee Dulyayangkul, Matthew B. Avison

**Affiliations:** School of Cellular & Molecular Medicine, University of Bristol, Bristol. UK

## Abstract

Using modified *Klebsiella pneumoniae* clinical isolates, we show that *ramR* plus *ompK36* mutation together with production of the V239G variant KPC-3 confirs resistance to ceftazidime/avibactam plus aztreonam, imipenem/relebactam and meropenem/vaborbactam, but not cefepime/taniborbactam. This is because the V239G variant does not generate collateral β-lactam susceptibility as do many other KPC-3 variants associated with ceftazidime/avibactam resistance. Additional mutation of *ompK35* and carriage of a plasmid expressing the OXA-48-like carbapenemase OXA-232 was required to confer cefepime/taniborbactam resistance.

---

Aztreonam/avibactam (AZT/AVI) is a β-lactam/β-lactamase inhibitor combination currently in clinical trials, which has activity against Enterobacterales producing metallo-carbapenemases and those with aztreonam-hydrolysing enzymes such as plasmid-mediated AmpCs (pAmpCs), extended-spectrum β-lactamases (ESBLs) and the serine carbapenemase KPC. All these enzymes are increasingly carried in *Klebsiella pneumoniae*, and yet few studies have been performed to consider mechanisms of AZT/AVI resistance in this species. It was recently reported that among 8787 Enterobacterales isolates, 17 were AZT-AVI resistant. Of these, three *Klebsiella* spp, were identified. Production of the pAmpC, DHA-1 plus *acrA* efflux pump gene overexpression and mutation of *ompK35* or *ompK36* porins were identified in two resistant isolates. The other produced the ESBL PER-2 and carried an *ompK35* loss of function mutation (1). In one *in vitro* study, selecting AZT/AVI resistance identified mutations in the pAmpC, CMY-16 in a *K. pneumoniae* strain (2). Avibactam is currently in clinical use partnered by ceftazidime (CAZ/AVI) and here, mutations in KPC are known to confer resistance. However, such mutations tend to reduce hydrolytic activity to β-lactams other than ceftazidime, including carbapenems and aztreonam (3-6). Accordingly, it is conceivable that such mutant KPC enzymes might not confer AZT/AVI resistance.

Another recently licenced β-lactam/β-lactamase inhibitor combination is imipenem/relebactam (IMI/REL). Unlike AZT/AVI, this does not have efficacy against isolates producing metello carbapenemases, but is generally efficacious against Enterobacterales producing pAmpC, KPC and ESBLs (7). Again, analysis of clinical isolates shows that IMI/REL resistance in *K. pneumoniae* is rare, but resistant isolates have mutations in or reduced expression of *ompK35* and/or *ompK36* porin genes and/or increased *acrA* efflux pump gene expression, alongside ESBL production (8). Similar impacts of porin and efflux pump production on IMI/REL susceptibility have been seen in *in vitro* studies using KPC-producing isolates (9).

Given seeming overlaps between AZT/AVI and IMI/REL resistance mechanisms in *K. pneumoniae*, we set out to dissect the mechanisms contributing to resistance to each in *K. pneumoniae* using a bank of clinical isolates and targeted recombinants having fully defined genotypes. **Table 1** reports MICs (determined using CLSI broth microdilution methodology [10,11]) of these combinations against a collection of clinical isolates, which have been previously described (12) and their β-lactam resistance genotypes characterised (13). All isolates, whether producing carbapenemases of classes A (KPC-3), B (NDM-1) or D (OXA-232) were AZT/AVI susceptible, but the NDM-1/OXA-232 producer KP4 was, as expected IMI/REL resistant, as was the OXA-232 producer KP11, though with lower MICs (**Table 1**). Notably, KP4 and KP11 have *ramR* mutations (12), which leads to over production of AcrAB-TolC efflux pump, and reduced production of the OmpK35 porin in *K. pneumoniae* (14). Nonetheless, a *ramR* mutant clinical isolate producing KPC-3, KP30, was susceptible to both AZT/AVI and IMI/REL (**Table 1**) so we conclude that modulating production of these permeability-associated proteins is not sufficient to give resistance to either β-lactam/β-lactamase inhibitor combination in a KPC-3 positive background.

**Table 1.** MICs of aztreonam and imipenem with or without avibactam or relebactam against *K. pneumoniae* clinical isolates

| Isolate (relevant genotype) | MIC ($\mu\text{g.ml}^{-1}$) | | | |
| --- | --- | --- | --- | --- |
|  | AZT | AZT/AVI | IMI | IMI/REL |
| KP31 (wild-type) | $\leq 0.5$ | $\leq 0.5$ | $\leq 0.5$ | $\leq 0.5$ |
| KP21 (*ramR* TEM-1) | $\leq 0.5$ | $\leq 0.5$ | $\leq 0.5$ | $\leq 0.5$ |
| KP11 (*ramR* OXA-232 CTX-M-15 TEM-1) | >128 | $\leq 0.5$ | 4 | 4 |
| KP30 (*ramR ompK35* KPC-3 TEM-1) | >128 | 1 | >128 | 1 |
| KP4 (*ramR* NDM-1 OXA-232 CTX-M-15 TEM-1) | >128 | $\leq 0.5$ | 64 | 16 |

Shading indicates resistance based on CLSI breakpoints (11)

To investigate the role of *bla*_KPC-3_ mutations known to be associated with CAZ/AVI resistance (15) in AZT/AVI and IMI/REL susceptibility, we took clinical *K. pneumoniae* isolate KP21, which is a *ramR* mutant and fully susceptible to AZT and IMI (**Table 1**). We introduced *bla*_KPC-3_ on a plasmid (pKPC-3), either wild-type or following site-directed mutagenesis to create a D178Y or V239G mutations (numbering based on the original KPC-3 sequence nomenclature [16]), previously associated with CAZ/AVI resistance (15). The construction of these plasmids has been reported previously (17). Reduced MICs of AZT and IMI were observed against KP21 carrying the D178Y variant, compared with KP21 carrying wild-type KPC-3 (**Table 2**). This phenomenon of reduced spectrum of β-lactamase activity has been described for other *bla*_KPC-3_ mutants associated with CAZ/AVI resistance (3-6). However, in a KP21 background, this reduction in activity was seen to a greater extent for the D178Y mutant, than the V239G variant (**Table 2**). This observation fits with previous reports that *K. pneumoniae* carrying the V239G mutant *bla*_KPC-3_ remain meropenem resistant, while those carrying the D178Y mutant are meropenem susceptible (15, 17). However, AZT/AVI and IMI/REL MICs were not greatly elevated against KP21 carrying pKPC-3 V239G in comparison with KP21 carrying pKPC-3, and all these KP21 recombinants remained AZT/AVI and IMI/REL susceptible (**Table 2**). We conclude, therefore, that mutating *bla*_KPC-3_ in a way that gives CAZ/AVI resistance is not sufficient to give AZT/AVI or IMI/REL resistance, even in a *ramR* mutant *K. pneumoniae* background.

**Table 2.** MICs of aztreonam or imipenem with or without avibactam or relebactam against derivatives of *K. pneumoniae* clinical isolates KP21 and KP47

| Isolate | MIC ($\mu\text{g.ml}^{-1}$) | | | |
| --- | --- | --- | --- | --- |
|  | AZT | AZT/AVI | IMI | IMI/REL |
| KP21[*ramR*] pUBYT | 0.5 | $\leq 0.5$ | 0.5 | $\leq 0.5$ |
| KP21[*ramR*] pKPC-3 | >128 | 1 | 64 | 2 |
| KP21[*ramR*] pKPC-3 D178Y | 1 | $\leq 0.5$ | 1 | 0.5 |
| KP21[*ramR*] pKPC-3 V239G | >128 | 2 | 32 | 2 |
| KP21[*ramR*] pUBYT pOXA-232 | 0.5 | $\leq 0.5$ | 2 | 1 |
| KP21[*ramR*] pKPC-3 pOXA-232 | >128 | 2 | 128 | 2 |
| KP21[*ramR*] pKPC-3 D178Y pOXA-232 | 16 | 2 | 2 | 2 |
| KP21[*ramR*] pKPC-3 V239G pOXA-232 | >128 | 4 | 32 | 8 |
| KP21[*ramR*] *ompK36* pUBYT | 0.5 | 1 | 2 | 0.5 |
| KP21[*ramR*] *ompK36* pKPC-3 | >128 | 2 | 128 | 2 |
| KP21[*ramR*] *ompK36* pKPC-3 D178Y | 8 | 2 | 1 | 0.5 |
| KP21[*ramR*] *ompK36* pKPC-3 V239G | >128 | 16 | 128 | 4 |
| KP21[*ramR*] *ompK36* pUBYT pOXA-232 | 1 | 1 | 4 | 2 |
| KP21[*ramR*] *ompK36* pKPC-3 pOXA-232 | >128 | 2 | >128 | 32 |
| KP21[*ramR*] *ompK36* pKPC-3 D178Y pOXA-232 | 8 | 4 | 8 | 2 |
| KP21[*ramR*] *ompK36* pKPC-3 V239G pOXA-232 | >128 | 16 | >128 | 32 |
| KP47 *ompK36* pUBYT | 0.5 | $\leq 0.5$ | 0.5 | 0.5 |
| KP47 *ompK36* pKPC-3 | >128 | 1 | >128 | 8 |
| KP47 *ompK36* pKPC-3 D178Y | 16 | 2 | 1 | 0.5 |
| KP47 *ompK36* pKPC-3 V239G | >128 | 4 | >128 | 16 |
| KP47 *ompK36* pUBYT pOXA-232 | 0.5 | $\leq 0.5$ | 2 | 1 |
| KP47 *ompK36* pKPC-3 pOXA-232 | >128 | 1 | >128 | 8 |
| KP47 *ompK36* pKPC-3 D178Y pOXA-232 | 16 | 2 | 4 | 2 |
| KP47 *ompK36* pKPC-3 V239G pOXA-232 | >128 | 4 | >128 | 16 |

Shading indicates resistance based on CLSI breakpoints (11)

Addition of an OXA-232 (class D carbapenemase) plasmid (pOXA-232, as described in our previous work [17]) to the KP21 recombinant carrying pKPC-3 V239G conferred IMI/REL (but not AZT/AVI) resistance, but this was not seen for KP21 recombinants carrying pKPC-3 D178Y or pKPC-3 (**Table 2**). Disruption of the *ompK36* porin gene in KP21 (as described previously[17]) conferred AZT/AVI and IMI/REL resistance when the recombinant was carrying pKPC-3 V239G, but not when it carried pKPC-3 D178Y or pKPC-3. Addition of pOXA-232 to the KP21 *ompK36* recombinants further raised MICs against the pKPC-3 V239G recombinant, and actually conferred IMI/REL resistance in the recombinant carrying pKPC-3, but not pKPC-3 D178Y (**Table 2**). Using the *ramR* wild-type isolate KP47 engineered to have an *ompK36* mutation we confirmed that *ramR* mutation is essential for the AZT/AVI resistance seen in KP21 *ompK36* pKPC-3 V239G, even following addition of pOXA-232 (**Table 2**).

We therefore conclude that three steps: mutation of *ramR*, mutation of *ompK36* and carriage of the V239G variant of *bla*_KPC-3_ is sufficient for *K. pneumoniae* to become resistant to both AZT/AVI and IMI/REL. However, prior to clinical approval of AZT/AVI, this combination is usually created clinically by adding AZT to CAZ/AVI therapy. A checkerboard assay confirmed that AZT/AVI and IMI/REL resistant derivative KP21[*ramR*] *ompK36* pKPC-3 V239G is also resistant to CAZ/AVI plus AZT, with MICs of CAZ (>32 µg.ml^-1^) and AZT (16 µg.ml^-1^) against this recombinant (**Figure 1**).

**Figure 1.**
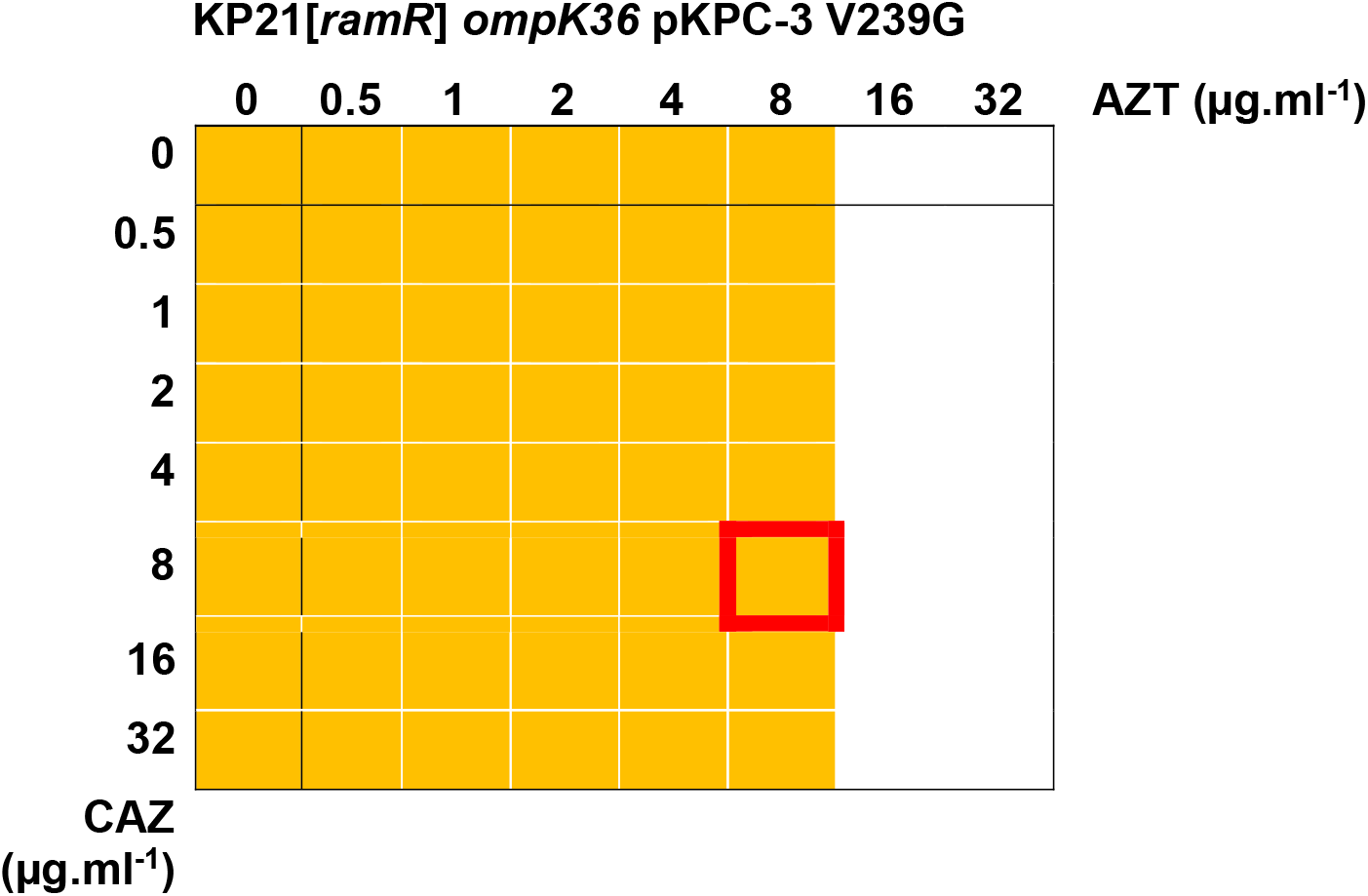
Checkerboard assays for CAZ and AZT in the presence of AVI against *K. pneumoniae* KP21[*ramR*] *ompK36* producing KPC-3 V239G. The image represents duplicate assays for an 8×8 array of wells in a 96-well plate. All wells contained CA-MHB including avibactam (4 µg.ml^-1^). A serial dilution of aztreonam (AZT, x-axis) and ceftazidime (CAZ, y-axis) was created from 32 µg.ml^-1^ in each plate as recorded. All wells were inoculated with a suspension of bacteria, made as per CLSI microtiter MIC guidelines (10), and the plate was incubated at 37°C for 20 h. Growth was recorded by measuring OD_600_ and growth above background (broth) is recorded as a yellow block; no growth is recorded as a white block. Growth in the red edged block indicates resistance to both AZT and CAZ.

We have previously shown that this combination of *ramR* and *ompK36* mutation coupled with acquisition of pKPC-3 V239G also gives resistance to another licenced β-lactam/β-lactamase inhibitor combination meropenem/vaborbactam (17). Finally, therefore, we considered the MIC of another combination in late stage clinical trials, cefepime/taniborbactam (18) against this derivative. Notably, in this *ramR ompK36* mutant background pKPC-3 D178Y supported lower cefepime MICs than pKPC-3 (**Table 3**), as seen for the other β-lactams (**Table 2**) and this was also true for cefepime/taniborbactam (**Table 3**). But again, the V239G mutant did not suffer from this reducton in cefepime MIC, and the MIC of cefepime/taniborbactam was the same against pKPC-3 and pKPC-3 V239G recombinants of KP21[*ramR*] *ompK36*, being 8 µg.ml^-1^, which is one doubling dilution below the cefepime resistance breakpoint (11) (**Table 3**). Further addition of pOXA-232 elevated cefepime MICs against the recombinants (**Table 3**), as expected since OXA enzymes are known to hydrolyse cefepime (19), but the cefepime/taniborbactam MIC remained at ≤ 8 µg.ml^-1^ indicating successful inhibition of OXA-232. However, additional insertional inactivation of *ompK35* porin gene (performed as described previously [17]) pushed the cefepime/taniborbactam MIC against KP21[*ramR*] *ompK36* recombinants carrying pOXA-232 and pKPC-3 V239G (but not pKPC-3 D178Y) to 16 µg.ml^-1^, which is classed as cefepime resistant (**Table 3**).

**Table 3.** MICs of cefepime/taniborbactam against derivatives of *K. pneumoniae* clinical isolate KP21

| Isolate | MIC ($\mu\text{g.ml}^{-1}$) | |
| --- | --- | --- |
|  | Cefepime | Cefepime/<br>Taniborbactam |
| KP21[*ramR*] *ompK36* pUBYT | 8 | 1 |
| KP21[*ramR*] *ompK36* pKPC-3 | >256 | 8 |
| KP21[*ramR*] *ompK36* pKPC-3 D178Y | 16 | 2 |
| KP21[*ramR*] *ompK36* pKPC-3 V239G | >256 | 8 |
| KP21[*ramR*] *ompK36* pUBYT pOXA-232 | 8 | 1 |
| KP21[*ramR*] *ompK36* pKPC-3 pOXA-232 | >256 | 8 |
| KP21[*ramR*] *ompK36* pKPC-3 D178Y pOXA-232 | 64 | 1 |
| KP21[*ramR*] *ompK36* pKPC-3 V239G pOXA-232 | >256 | 8 |
| KP21[*ramR*] *ompK36 ompK35* pUBYT | 8 | 1 |
| KP21[*ramR*] *ompK36 ompK35* pKPC-3 | >256 | 8 |
| KP21[*ramR*] *ompK36 ompK35* pKPC-3 D178Y | 16 | 2 |
| KP21[*ramR*] *ompK36 ompK35* pKPC-3 V239G | >256 | 8 |
| KP21[*ramR*] *ompK36 ompK35* pUBYT pOXA-232 | 8 | 2 |
| KP21[*ramR*] *ompK36 ompK35* pKPC-3 pOXA-232 | >256 | 8 |
| KP21[*ramR*] *ompK36 ompK35* pKPC-3 D178Y pOXA-232 | 64 | 2 |
| KP21[*ramR*] *ompK36 ompK35* pKPC-3 V239G pOXA-232 | >256 | 16 |

We conclude, therefore, that whilst three events (*ramR, ompK36, bla*_KPC-3_ V239G) are sufficient to cause AZT/AVI, CAZ/AVI/AZT, IMI/REL and meropenem/vaborbactam resistance in *K. pneumoniae*, additional events are required to give cefepime/taniborbactam resistance. Furthermore, whilst many *bla*_KPC-3_ mutations leading to CAZ/AVI resistance do come with the collateral effect of increased susceptibility to carbapenems, late generation cephalosporins and AZT, KPC-3 V239G does not suffer from this effect to the same degree. This explains why KPC-3 V239G, rather than KPC-3 D178Y, which does suffer from collateral increased susceptibility is able to confer resistance to multiple β-lactam/β-lactamase inhibitor combinations, provided their accumulation is slowed. The implication here is that KPC-3 V239G is inhibited to a lesser extent than KPC-3, rather than having a more ceftazidime focussed hydrolytic spectrum, as does KPC-3 D178Y (20). Accordingly, the emergenece of this *bla*_KPC-3_ variant should be watched with caution.

## Transparency declaration

The authors declare no conflict of interests.

